# Multi-Head Attention-based U-Nets for Predicting Protein Domain Boundaries Using 1D Sequence Features and 2D Distance Maps

**DOI:** 10.1101/2022.04.08.487689

**Authors:** Sajid Mahmud, Zhiye Guo, Farhan Quadir, Jian Liu, Jianlin Cheng

## Abstract

The information about the domain architecture of proteins is useful for studying protein structure and function. However, accurate prediction of protein domain boundaries (i.e., sequence regions separating two domains) from sequence remains a significant challenge. In this work, we develop a deep learning method based on multi-head U-Nets (called DistDom) to predict protein domain boundaries utilizing 1D sequence features and predicted 2D inter-residue distance map as input. The 1D features contain the evolutionary and physicochemical information of protein sequences, whereas the 2D distance map includes the structural information of proteins that was rarely used in domain boundary prediction before. The 1D and 2D features are processed by the 1D and 2D U-Nets respectively to generate hidden features. The hidden features are then used by the multi-head attention to predict the probability of each residue of a protein being in a domain boundary, leveraging both local and global information in the features. The residue-level domain boundary predictions can be used to classify proteins as single-domain or multi-domain proteins. It classifies the CASP14 single-domain and multi-domain targets at the accuracy of 69.1%, 2.67% more accurate than the state-of-the-art method. Tested on the CASP14 multi-domain protein targets with expert annotated domain boundaries, the average per-target F1 measure score of the domain boundary prediction by DistDom is 0.263, 29.56% higher than the state-of-the-art method.

## Introduction

Protein domains are cohesive structural regions of a protein chain, which usually assume a compact structure. Domains are the structural, functional or evolutionary units of proteins [1]. They also act as the building blocks of larger proteins [2]. The information about the location of domains is useful for improving protein structure prediction [3] and analyzing protein function. Thus, identification of protein domain boundaries is important for addressing the challenge in protein structure and function annotations [4] as well as exploring the protein structure and function space [5]. As a large amount of protein sequences is being produced on a daily basis, there is a significant need of automated methods for predicting domain boundaries in protein sequences [6].

There are two types of approaches for predicting protein domain boundaries from sequences without known experimental structures, i.e., template-based and ab initio prediction. In the case of template-based prediction, the goal is to find homologous template sequences that have known domain information for a target protein. These alignments between the templates and a target are used to transfer the domain boundary information from the homologous sequences to the target protein. Some template-based methods are CHOP [7], DomPred [8], SSEP-Domain [9], ThreaDom [10], CLADE [11], MetaCLADE [12] and SnapDRAGON [13]. These methods can be effective and accurate when close templates are found. However, if no highly similar templates with known domain information are found, the performance of these models declines sharply [1]. The ab initio prediction can be broadly applied to any protein as it does not require a homologous template with known domain information. The ab initio methods are mostly based on statistical and machine learning models, which are trained to predict domain boundaries from sequences and sequence-derived features. Some ab initio methods are CHOPnet [7], PPRODO [14], DOMpro [15], KemaDom [16], DomainDiscovery [17], IGRN [18], DomSVR [19], DoBo [6], DROP [20], DomHR [21], PDP-CON [22], ConDo [23], DNN-Dom [24], DeepDom [1], FuPred [25]. As ab initio methods do not have similar templates as a guidance for domain boundary prediction, their performance tends to be less accurate than the template-based methods for targets having templates available. This is partly because extracting useful information from sequences for boundary prediction is still an open, challenging problem [1]. The expert-curated features through a manual, laborious process used by the existing methods are not sufficiently informative for accurate domain prediction. Deep learning is a powerful technique to automatically generate features from raw input to improve prediction accuracy [26, 27]. In this work, we develop U-Net deep learning architectures [28] that achieved success in image segmentation to automatically extract the features from 1D and 2D input information generated from protein sequences to predict domain boundaries. 1D sequential information such as sequence profiles used in secondary structure prediction [29] as well as 2D residue-residue distance map predicted from sequences [30] is used as input. Because domain boundaries are global properties of a protein that may depend on residues far away, we integrate a multi-head attention mechanism approach [31] with U-Nets to extract long-range information from the 1D and 2D inputs to predict whether a residue falls into a domain boundary or not. The attention mechanism can capture signals related to domain boundaries occurring anywhere in the input more effectively than traditional recurrent neural networks [32] or more advanced long-and short-term memory (LSTM) networks [33]. Because the predicted 2D distance map is a rather unique feature of our method that was rarely used by the existing methods, our method is called DistDom. In addition to the novel multi-head attention based 1D and 2D U-Net architectures to integrate 2D distance maps with 1D sequence information to predict domain boundaries, DistDom substantially improves the prediction accuracy over the existing methods.

## Methods

### Training, Validation and Test Datasets

We use 4072 protein targets from the Topdomain dataset [34] to create the training and validation data. These proteins were split 80% for training and 20% for validation. We also collected the targets with domain information curated by CASP organizers from the archive of CASP7-14 experiments [35] as independent test data. 619 proteins (193 multi domain targets, 426 single domain targets) from the CASP7-13 are used as one test data and 55 proteins (20 multi domain targets, 35 single domain) from CASP14 are used as another test dataset. The domains of the CASP14 proteins have been manually annotated by CASP14 organizers and assessors and therefore are of high quality. To rigorously evaluate the prediction performance, we remove the proteins in the validation test data sets that have more than 30% sequence identity with the training dataset according to PSI-BLAST [36] alignments. The final Topdomain validation set has 793 protein targets, the final CASP7-13 test dataset has 610 targets, and the CASP14 dataset has 55 targets. Because a domain boundary region usually spans a number of residues between two domains, 10 residues within an expert-annotated boundary position on the sequence of a target are considered boundary residues 1 (i.e., positive examples). All other residues are considered non-boundary ones (i.e., negative examples). In addition, a region separated by a domain boundary must have at least 40 residues to be considered a separate domain. After labeling the residues, the domain boundary prediction problem is to predict if a residue is a boundary residue or not.

### Input Feature Generation

We generate two kinds of input features including two-dimensional (2D) residue-residue distance maps and one-dimensional (1D) sequential information for each protein, which are described below.

### 2D Distance Map

Residue-residue distance map is a concise representation for protein conformation, which can be used to reconstruct 3D protein structure. The distance map can be rather accurately predicted for many proteins from sequences [37, 38]. Therefore, predicted distance maps contain relevant information that can be used to predict protein domain boundaries. However, even though true residue-residue contact maps (i.e., binary distance maps) have been used to delineate protein domain boundaries, predicted distance maps have not been well used as input for deep learning methods to predict domain boundaries. In this work, we use DeepDist [30], an ab initio deep learning method for predicting residue-residue distance maps as input. For a protein sequence of length L, a distance map of L x L matrix contains the predicted Euclidean distance (d) between any two residues (i, j) in terms of angstrom. Because generally only distances smaller than a threshold contain useful information regarding protein structure, we use a distance threshold (e.g., 8,12,16 Å) to select all the predicted distances equal to or less than the threshold as input, while replacing all other distances with a special value (−1). The entire input matrix is then normalized by the min-max normalization to facilitate the training of deep learning models. Because the maximum value in the matrix is the threshold (t) and the minimum value is −1, a distance (d) in the matrix is normalized by the formula: (d - min) / (max - min) = (d + 1) / (t + 1). Three commonly used distance thresholds (i.e., 8,12,16 Å) are tested to select the best threshold. The shape of the 2D input is L x L. Figure 1 compares an original distance map and its normalized counterpart, demonstrating that the signals in the normalized map are more clearer than in the original one.

**Figure 1:**
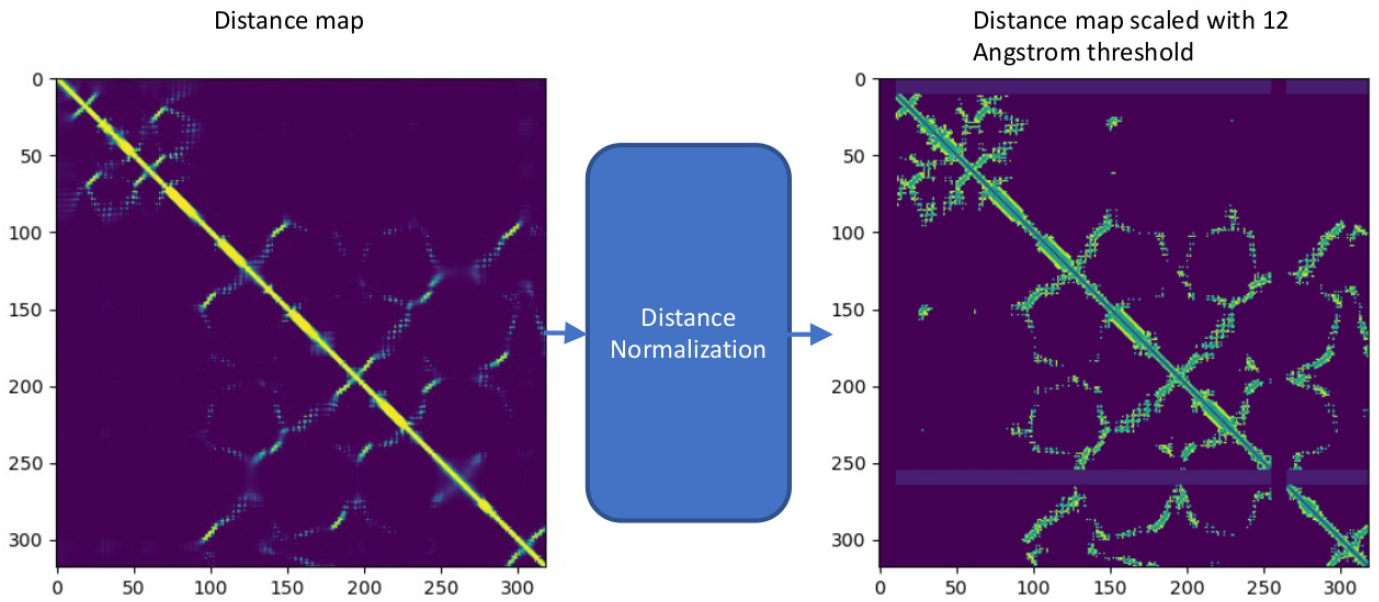
An example of distance map normalization for target T1101 from CASP14 dataset.

### 1D Sequential Information

The domain boundary prediction is a 1D prediction problem that has some similarity with other 1D structural feature prediction problems such as secondary structure prediction, in which sequence profile and residue conservation information are useful. Therefore, we use the input generation module of an accurate secondary structure predictor - DNSS2 [29] to generate 1D sequential input for the domain boundary prediction. For each position in a protein sequence of length L, the inputs include 21 numbers from the position specific scoring matrix (PSSM) generated by using PSI-BLAST to search the sequence against the UniRef90 sequence database, 20 emission probabilities and 7 transition probabilities extracted from the Hidden Markov Model (HMM) profile generated by using HHblits tool to search the uniclust30 database, 20 probabilities of 20 standard amino acids at the position calculated from the multiple sequence alignments (MSA) generated by using HHblits tool to search uniclust30 database, and 5 numbers derived from Atchley’s factor that describe physiochemical property of the residue at the position [29]. In total, there are 73 input numbers for each residue in a protein sequence. Therefore, the shape of the 1D input is L x 73, where L is the length of the sequence.

### Multi-head Attention-based U-Nets for Domain Boundary Prediction

The overall workflow of DistDom is depicted in Figure 2. The 2D input of shape *L* × *L* × 1 and 1D input of shape L x 73 are processed by 2D U-Net and 1D U-Net simultaneously and respectively. The 2D output of the 2D U-Net is put through an adaptive max pooling to reduce the dimension to *L* × 1. The output of the 1D U-Net has the shape of *L* × 1. The two outputs are concatenated together as the hidden features of shape of *L* × 2, which are used as the input for the multi-head attention layer to predict the probability that each residue is in a domain boundary or not. The shape of the final output is *L ×* 2, containing the probability of being in a domain boundary and not in a boundary.

**Figure 2:**
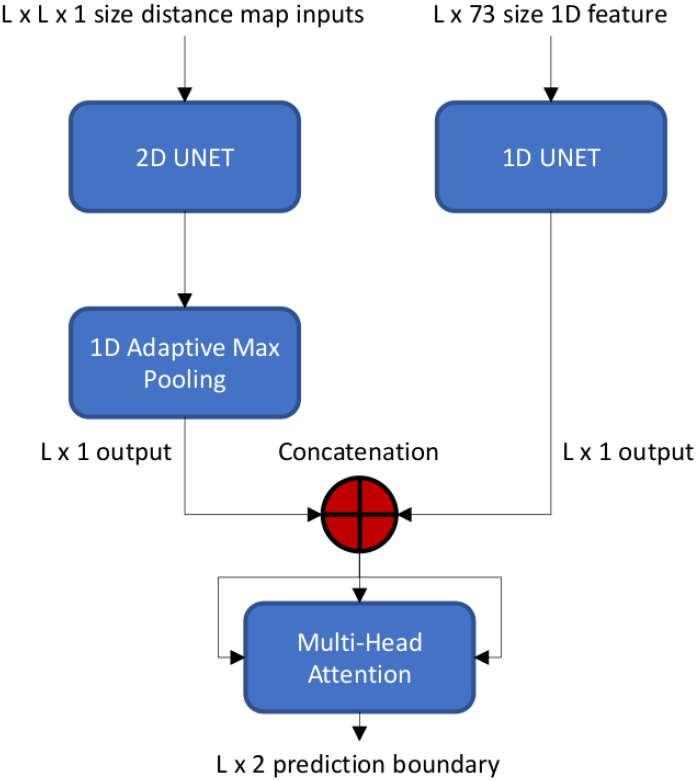
The overall architecture of DistDom. The 2D distance maps and 1D features are processed by a 2D U-Net and 1D U-Net respectively. The 2D U-Net output is converted by the 1D adaptive max pooling to the output of *L ×* 1. The *L ×* 1 outputs from the 1D U-Net and 2D-U-Net are combined together as the hidden features for the multi-head attention layer to predict the probability of being in a domain boundary or not.

### 2D U-Net

A customized 2D U-Net architecture [28] (Figure 3) is used to process the 2D distance map input. One difference between the customized 2D U-Net and the canonical 2D-UNet is that the former is designed to handle the distance map of variable dimension, but the latter is applied to the image of fixed dimension. It consists of three main components: downsampling component, upsampling component, and direct connections from the blocks in the downsampling component to their counterparts in the upsampling component. In the downsampling component, two consecutive convolutional layers form a block. Each 2D convolution layer consists of 2D convolution with a 3 × 3 kernel, 2D batch normalization layer and a ReLu activation function. The filter size of the initial convolutional layer is 64, whereas subsequent layers have filter size of 128, 256, 512 and 1024 respectively. The first block converts the 2D input (i.e., the normalized distance map) into a hidden feature map of *L × L ×* 64. The dimension of the feature map is reduced by half to *L/*2 × *L/*2 × 64 by the max pooling, which is processed by the next block to generate the hidden feature of the same dimension but doubled depth (i.e., *L/*2 × *L/*2 × 128). This process is repeated until a feature map with the shape of *L/*16 × *L/*16 × 1024 is generated. This reduced map is the latent embedded representation of the original distance map input. The upsampling process is a reverse process of the downsampling process by gradually reconstructing an output of the same dimension as that of the initial input from the final embedded hidden feature map generated by the downsampling process. Firstly, that embedded feature map is converted by a two-layer convolutional block to a L/16 x L/16 × 1024 feature map, which is then upsampled using bilinear interpolation to a feature map of *L/*8 *× L/*8 *×* 1024. This map is then concatenated by depth with its counterpart of the same dimension (*L/*8 *× L/*8 *×* 512) in the downsampling process through a direct connection [28] to produce a feature map of size *L/*8 *× L/*8 *×* 1536. This direct connection can account for some lost information in the downsampling process and also speed up learning like the widely used residual connection. Afterwards, the *L/*8 *× L/*8 *×* 1536 feature map goes through convolutional layers to produce a feature map of size *L/*8 *× L/*8 *×* 512. This process is repeated until the dimension of the feature map is returned to *L × L ×* 64, which is used to generate a final feature map of *L × L ×* 1 using 1 *×* 1 convolution.

**Figure 3:**
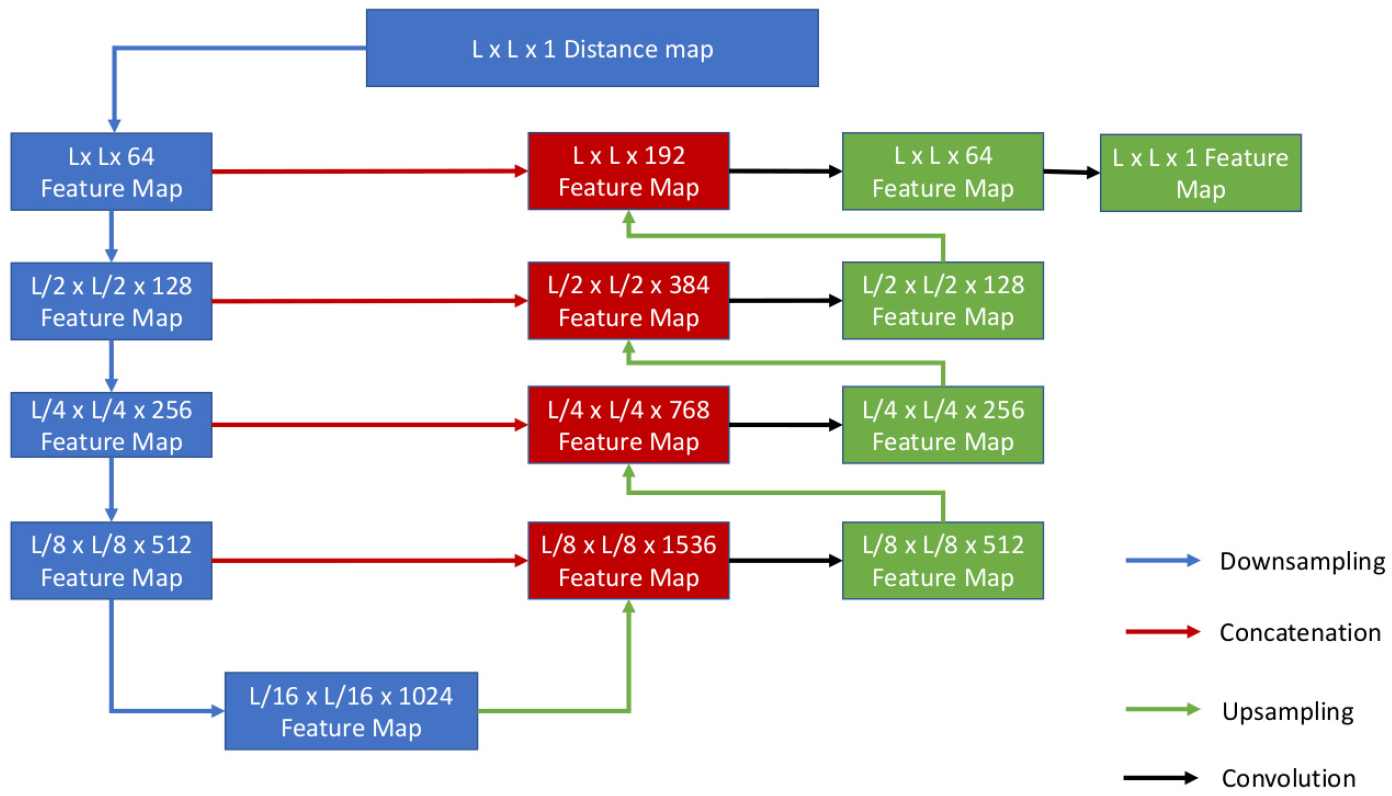
The 2D U-Net of processing 2D inputs. Adapted from the original U-Net paper [28]

### 1D U-Net

We generalize the U-Net typically applied to 2D input to a customized 1D U-Net architecture for processing the 1D input. The key difference between 1D U-Net and 2D U-Net is that it uses 1D convolution instead of the 2D convolution, and for upsampling, it uses 1D transposed convolution. The detailed structure of the 1D U-Net is depicted in Figure 4. It takes the 1D input of shape L x 73 to generate a L x 1 hidden feature map.

**Figure 4:**
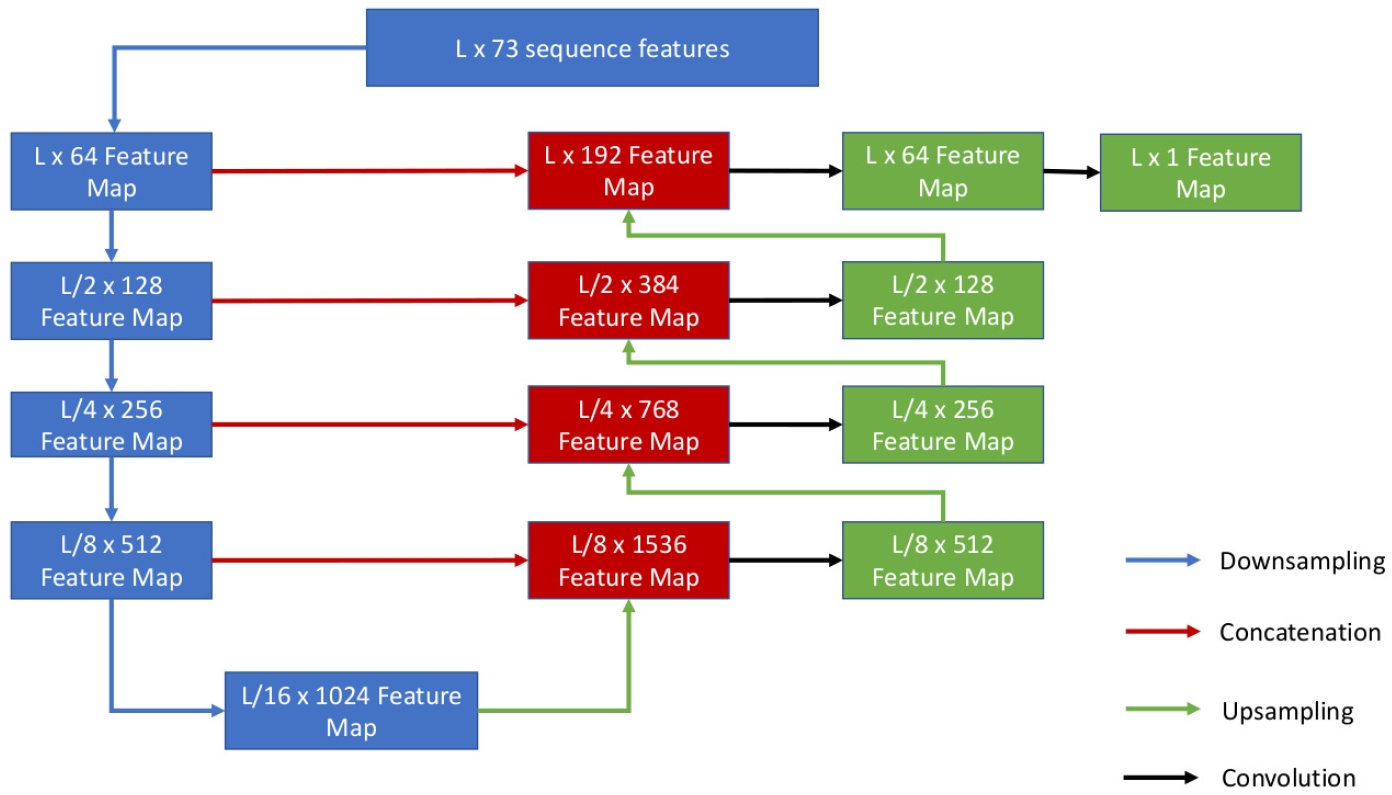
1D U-Net. The input of shape L x 73 is processed by the downsampling and upsampling process to generate a final hidden feature map of shape L x 1.

**Figure 5:**
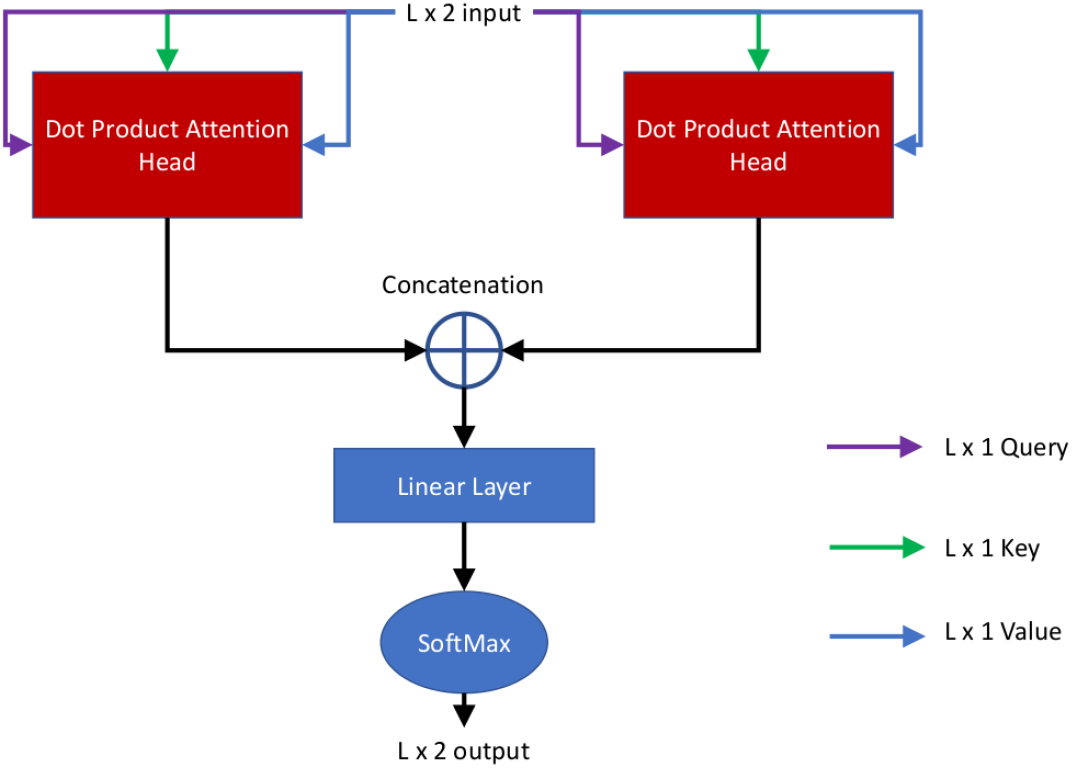
The multi-head attention mechanism. Here the *L ×* 2 input is split into two L x 1 inputs. Each of these L x 1 inputs are fed into a scaled dot product attention head as the query, key and values. There are two attention heads, each of those yielding an output of L x 1 size. The attention head outputs are concatenated to form a tensor of the size of *L ×* 2, which is passed through a linear layer. The matrix multiplication with a weight matrix is applied to the input in the linear layer to generate the activation for the softmax function to predict the probability of two classes (in a domain boundary or not). The final output is *L ×* 2, where L is the sequence length.

### Multi-Head Attention

The output feature map of the 2D U-Net branch and 1D U-Net branch above are concatenated to generate a feature map of *L ×* 2 size, which is used as the input for the following self-attention layer [31, 39] to predict if a position i is in a domain boundary. The attention mechanism uses a scaled dot product involving three quantities: query Q, key K, and value V for each input node to calculate the attention weight according to Equation 1.

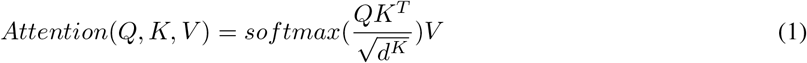

Here, *d*^*K*^ is the dimension of the K matrix for each attention head. In our case it is 1 for each attention head. In the self-attention process, the *Q, K* and *V* values are the same as the values of the concatenated hidden feature map from the two U-Nets. That is, the concatenated output of the 2D U-Net branch and 1D U-Net branch are each used as the query, key and value for each of their attention heads. The multi-head attention is applied to the concatenated *L ×* 2 feature map, which is split into two *L ×* 1 sub feature maps. One head is applied to the *L ×* 1 sub-feature map generated from the 1D input features and another to the *L ×* 1 sub-feature map generated from the 2D input features. The outputs of the two attention heads are concatenated as input of *L ×* 2, which is used by a linear layer to generate an output of size *L ×* 2 by multiplying the input by a weight matrix of size 2 × 2. Here 2 is the number of heads in the multi-head attention. Each row (i) in *L ×* 2 output from the linear layer is used by the softmax function to predict if residue i is in a domain boundary or not. There are two predicted probabilities (in a boundary and not in a boundary) for each residue whose sum is equal to 1. The multi-head attentive U-Nets can handle protein sequences of variable length because the same convolution and attention operations can be applied to protein sequences of any length. Therefore, no reshaping of the input data to a fixed length is needed, avoiding any potential pitfalls and biases posed by it.

### Training and Validation

We use the GPU computing resource on the Summit supercomputer provided by Oak Ridge National Laboratory (ORNL) to train the deep learning network above. The Summit cluster [40, 41] provides many compute nodes each having 6 GPUs and 16 GB of memory, which enables the distributed deep learning training. We train 2D U-Net, 1D U-Net, and the multi-head attention layer on three separate GPU nodes. The training is done with Adam optimizer [42] with a learning rate of 0.0001. The loss function is the cross-entropy [43]. Because only a small portion of residues are boundary residues (positive examples), we also test the weighted entropy loss. The weights for the positive and negative examples are 10 and 1, respectively. The early stopping is used to stop the training when the average per-target F1 measure score (2 x precision x recall / (precision + recall)) on the validation data does not increase. The best-performing model on the validation data is saved for testing.

### Code Availability

The code for predicting domain boundaries is available at the GitHub repository https://github.com/jianlin-cheng/DistDom.git.

## Results and Discussion

DistDom is trained on Topdomain [34] training dataset and validated on Topdomain validation dataset. After the training and validation, it is tested on CASP7-13 and CASP14 [44] test datasets.

### Impact of distance thresholds on the prediction performance

DistDom is trained with three different distance threshold values (8, 12 and 16 Å) commonly used to reconstruct tertiary structures from distance maps to select predicted residue-residue distances as input for domain prediction. The average per target F1 scores on the validation and test datasets on multi-domain targets for the three thresholds are shown in Table 1. On all three datasets, 12 Å threshold leads the way in terms of the performance based on the average per target F1 score. Considering the results on all the datasets, 12 angstrom is chosen as the final distance threshold for all the experiments. It is worth noting that the average per target F1 score reported in Table 1 is the average of F1 scores calculated for each target. The average per target F1 score is not a geometric mean of the average per target precision and average per target recall. Instead, the precision, recall and F1 score for each target is calculated individually, and the F1 scores of all the targets are averaged to produce the average per target F1 score.

**Table 1:** The average per target F1 score of using different distance thresholds (8, 12 and 16 Å) for domain boundary prediction on the multi-domain proteins in the three datasets: (1) Topdomain validation data; (2) CASP7-13 test data; and (3) CASP14 test data. Bold font denotes the best result.

| Dataset | 8Å | 12Å | 16Å |
| --- | --- | --- | --- |
| Topdomain Validation | 0.457 | 0.470 | 0.268 |
| CASP7-13 | 0.244 | 0.291 | 0.134 |
| CASP14 | 0.232 | 0.263 | 0.201 |

### Selection of cut-off decision threshold for domain boundary prediction

To decide a cut-off probability threshold needed to convert the predicted domain boundary probabilities to binary predictions (in a domain boundary or not), we test different cut-off thresholds between 0 and 1, with an interval of 0.01 on the Topdomain validation dataset to find the threshold yielding the highest F1 score. We plot the average per target precision, recall and F1 scores against the cut off values to choose a cut-off threshold (Figure 6).

**Figure 6:**
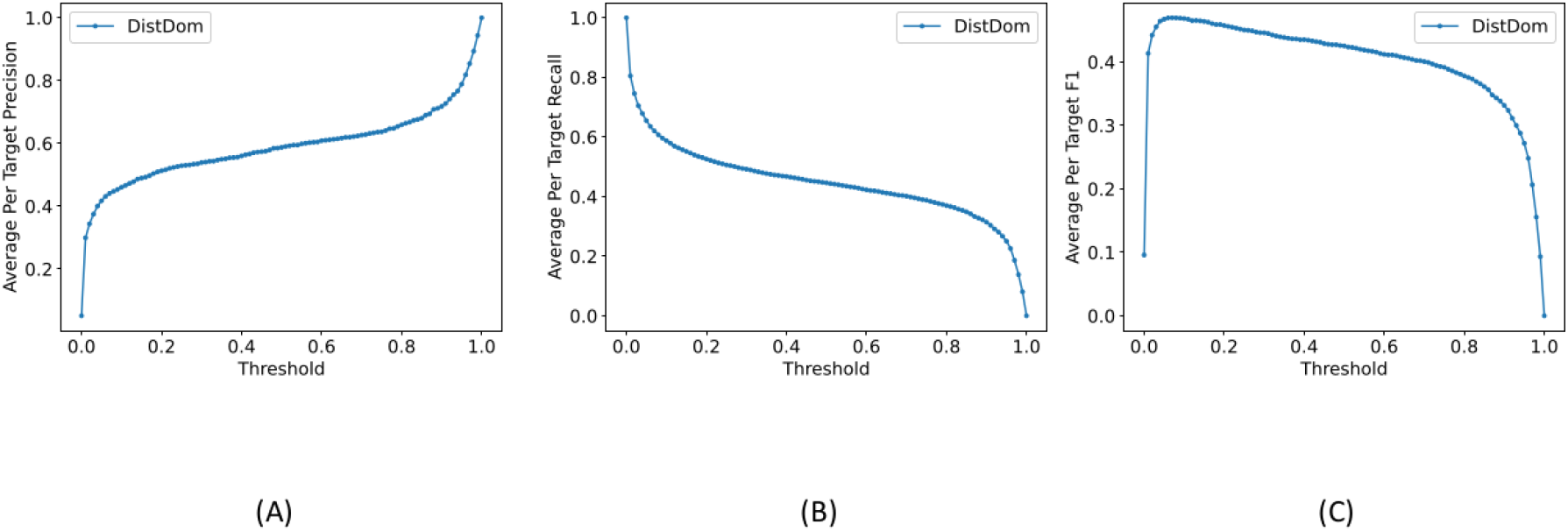
Comparison of average per target F1 scores on the Topdomain validation dataset with and without the distance normalization for multi domain targets.

### Impact of distance normalization on the prediction performance

To study the impact of the distance normalization, we compare the performance of DistDom trained with the distance normalization at 12 Å distance threshold and without distance normalization. The results on the validation dataset are reported in Table 2. The DistDom with the distance normalization outperforms the model without distance normalization in every aspect. For instance, the average per-target F1-score of using the normalization on all the multi domain targets is 0.470, higher than 0.377 without it.

**Table 2:** Comparison of average per target F1 scores on the Topdomain validation dataset with and without the distance normalization for multi domain targets.

| No Normalization | Normalization |
| --- | --- |
| 0.377 | 0.470 |

### Impact of 1D and 2D inputs on the prediction performance

We perform an ablation study to measure the effectiveness of 1D and 2D inputs. We conduct three experiments: one with 1D input only, one with only the 2D input, and one combining both inputs together. The results of the three experiments on the multi-domain proteins in the Topdomain validation dataset, CASP7-13 test dataset, and CASP14 test dataset are reported in Table 3. The results in Table 3 show that the combination of 1D and 2D inputs together consistently performs better than using either the 1D or 2D input only. The average per target F1 scores reported for 1D and 2D combined (0.470 on Topdomain validation, 0.291 on CASP7-13 and 0.263 on CASP14 dataset) are all higher than the average per target F1 scores for using 1D or 2D input alone.

**Table 3:** Comparison of the average per target F1 scores of using 1D input only, 2D input only and both 1D and 2D.

| Dataset | 1D Input Only | 2D Input Only | 1D and 2D combined (DistDom) |
| --- | --- | --- | --- |
| Topdomain Validation | 0.291 | 0.383 | 0.470 |
| CASP7-13 | 0.153 | 0.285 | 0.291 |
| CASP14 | 0.112 | 0.203 | 0.263 |

### Comparison of weighted and unweighted loss functions

Because there are many more negative examples than positive examples, we test a weighted cross-entropy loss function. The 1:10 weight ratio is applied to negative (non-boundary) class and positive (boundary) class, respectively. Table 4 compares the F1 score of using the weighted loss function and the unweighted one on the multi-domain proteins in the Topdomian validation dataset. The unweighted loss function works better than the weighted loss function overall. Therefore, the unweighted loss function is used to train the final DistDom model.

**Table 4:** Results of the unweighted and weighted loss functions on Topdomain validation dataset Model Average per target F1 score on multi-domain proteins

| Model | Average per target F1 score on multi-domain proteins |
| --- | --- |
| DistDom (unweighted) | 0.470 |
| DistDom (weighted) | 0.402 |

### Comparison with existing methods on the CASP14 test dataset

We compare DistDom with two other most recent methods - DeepDom and FUPred on the multi-domain targets in the CASP14 dataset. DeepDom [1] is based on a bidirectional LSTM network. FUPred [25] is a method that utilizes FUScore in an ad-hoc manner with deep learning-predicted contact maps to predict the location of domain boundaries. As the DeepDom predicts the probability of each residue belonging to the boundary or non-boundary, we use its default decision threshold (0.42) to convert the predicted probabilities into binary predictions. The same evaluation procedure used in DeepDom is used to evaluate it here. Table 5 reports the average per-target F1 scores of domain boundary prediction on the CASP14 multi-domain targets. The results show that DistDom performs better than DeepDom and FUPred on both single-domain and multi-domain targets. For instance, the average per-target F1 score on all the CASP14 multi-domain targets is 0.263, which is higher than 0.203 of FUPred and 0.134 of DeepDom.

**Table 5:** Results of DistDom, DeepDom, and FUPred on the multi-domain targets in the CASP14 dataset in terms of Average Per Target F1 Score

| Method | Average per target F1 score |
| --- | --- |
| DistDom | 0.263 |
| DeepDom | 0.134 |
| FUPred | 0.203 |

### Applying domain boundary prediction to distinguish multi-domain proteins from single-domain ones

A classification problem related to the protein domain boundary prediction is to classify if a protein is a multi-domain protein or a single-domain one. To classify the protein, DistDom is applied to predict the boundary residues of the protein. If there are *≥*10 residues predicted as boundary residues, the protein is classified as multi-domain, otherwise single-domain. The classification accuracy and MCC (Matthew’s Correlation Coefficient) are used to evaluate the classification results as in [25], which are defined as follows.

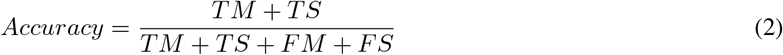

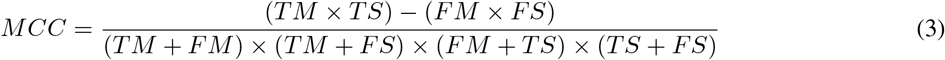

TM, TS, FM, and FS are the number of correct multi-domain prediction, number of correct single-domain predictions, number of false multi-domain predictions, and number of false single-domain predictions. The range of the Matthew’s Correlation Coefficient (MCC) is from −1 to 1.

The results of DistDom, FUPred, and DeepDom are reported in Table 6. DistDom performs best in terms of both the accuracy and MCC. The accuracy and MCC of DistDom are 0.691 and 0.339, which are 2.67% and 3.67% higher than FUPred.

**Table 6:**
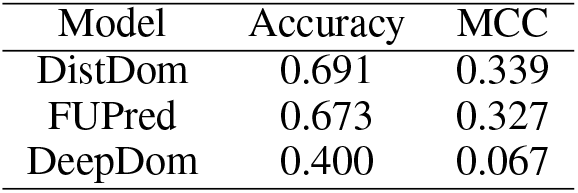
The accuracy and Matthew’s correlation coefficient of DistDom, FUPred, and DeepDom of classifying single-domain and multi-domain proteins in the CASP14 dataset.

| Model | Accuracy | MCC |
| --- | --- | --- |
| DistDom | 0.691 | 0.339 |
| FUPred | 0.673 | 0.327 |
| DeepDom | 0.400 | 0.067 |

### A good prediction example

Figure 7 illustrates a prediction example (PDB ID 5BNC (chain A) from Topdomain validation dataset). The predicted domain boundary overlaps well with the true domain boundary of his target. The precision of the prediction is 0.667, the recall is 1.0 and the F1 score is 0.800.

**Figure 7:**
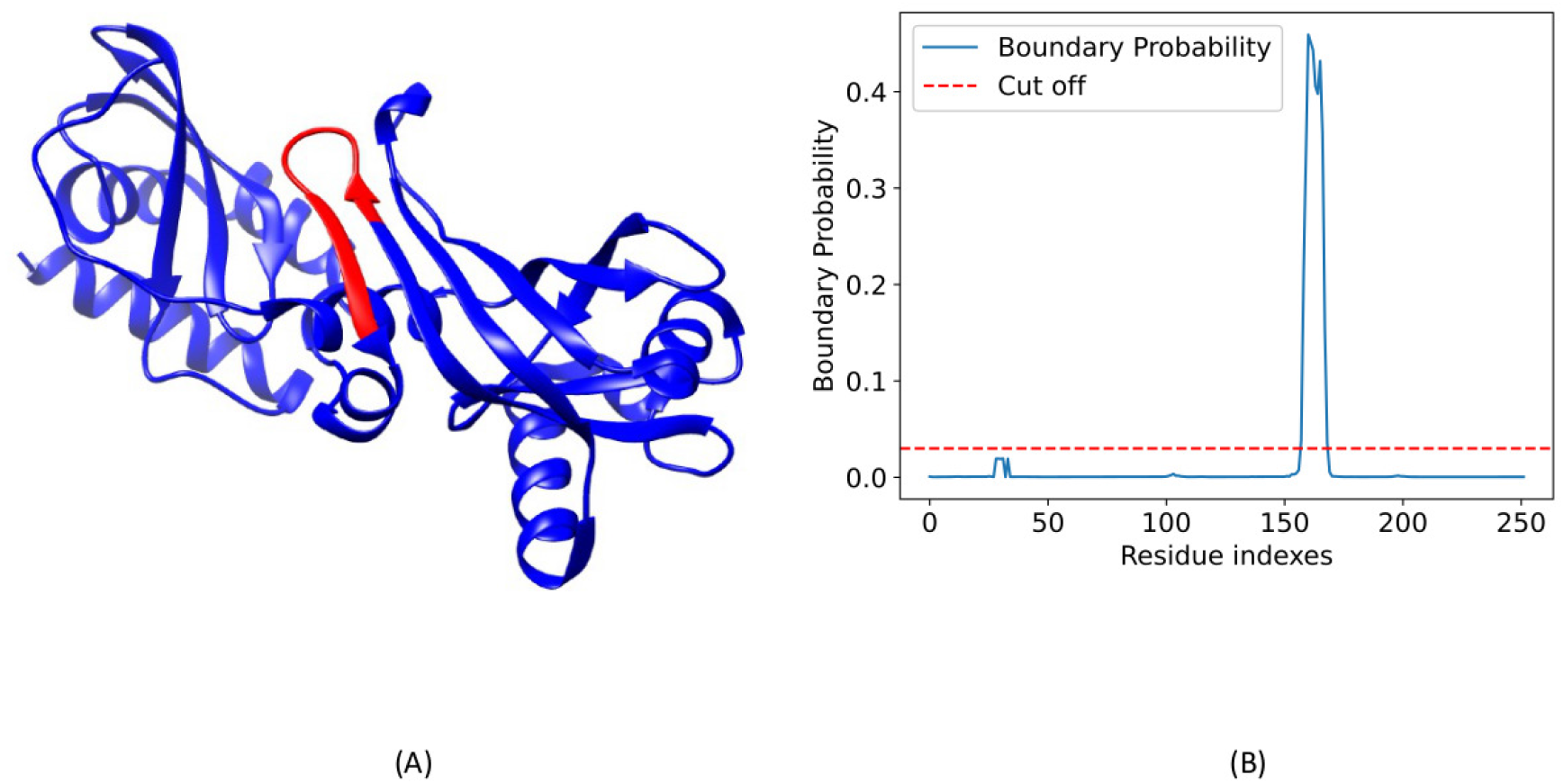
Domain boundary prediction for a protein target (PDB code 5BNC, chain A). The sub-figure on the left visualizes the true structure and the labeled true boundary (residues 158-167) in red color. The sub-figure on the right shows a plot of the predicted domain boundary probabilities against the residue positions. A red horizontal line shows the cut-off threshold (0.03). Residues 159-169 whose predicted probabilities are above 0.03 are the predicted boundary residues. The predicted boundary overlaps with the true boundary well.

## Conclusion

We develop an end-to-end deep learning based method based on the U-Nets and multi-head attention to improve the domain boundary prediction. It is the first method of combining traditional 1D sequence features and new 2D distance maps to predict protein domain boundaries. The 1D and 2D input features are processed by a customized 1D and 2D U-Net simultaneously to generate hidden features for the multi-head attention to predict the domain boundary. The experiment shows that, with the appropriate preprocessing (thresholding and normalization), the 2D distance map can be effectively used to predict protein domain boundaries. Combining 1D and 2D features via the multi-head attention can further improve the prediction accuracy. Moreover, the domain boundary predictions can be effectively used to classify if a protein is a single-domain protein or multi-domain one.

## Acknowledgements

Research reported in this publication was supported in part by Department of Energy grants (DE-AR0001213, DESC0020400, and DE-SC0021303), two NSF grants (DBI1759934 and IIS1763246), and an NIH grant (R01GM093123). This research used resources of the Oak Ridge Leadership Computing Facility at the Oak Ridge National Laboratory, which is supported by the Office of Science of the U.S. Department of Energy under Contract No. DE-AC05-00OR22725.

